# Mining impactful discoveries from the biomedical literature

**DOI:** 10.1101/2022.10.28.514184

**Authors:** Erwan Moreau, Orla Hardiman, Mark Heverin, Declan O’Sullivan

## Abstract

**Motivation:** Literature-Based Discovery (LBD) aims to help researchers to identify relations between concepts which are worthy of further investigation by text-mining the biomedical literature. While the LBD literature is rich and the field is considered mature, standard practice in the evaluation of LBD methods is methodologically poor and has not progressed on par with the domain. The lack of properly designed and decent-sized benchmark dataset hinders the progress of the field and its development into applications usable by biomedical experts.

**Results:** This work presents a method for mining past discoveries from the biomedical literature. It leverages the impact made by a discovery, using descriptive statistics to detect surges in the prevalence of a relation across time. This method allows the collection of a large amount of time-stamped discoveries which can be used for LBD evaluation or other applications. The validity of the method is tested against a baseline representing the state of the art “time sliced” method.

**Availability:** The source data used in this article are publicly available. The implementation and the resulting data are published under open-source license: https://github.com/erwanm/medline-discoveries (code) https://zenodo.org/record/5888572 (datasets). An online exploration tool is also provided at https://brainmend.adaptcentre.ie/.

## 1 Introduction

Nowadays virtually any biomedical research work is available almost instantly in digital form, but exploring the literature is made challenging by the ever-increasing amount of publications. Literature-Based Discovery (LBD) aims to automatically extract new insights from the scientific literature (Henry and McInnes 2017). Thus LBD is intended to assist researchers in identifying potentially interesting relations between concepts,^1^ potentially contributing to faster and broader scientific progress.

Swanson 1986a introduced LBD by linking dietary fish oil and Raynaud’s syndrome, noting that both concepts have a known relation to blood circulation. This promising first discovery was quickly followed by another one linking migraine and magnesium (Swanson 1988). These two initial discoveries are the most commonly used as benchmark for the evaluation of LBD systems (Thilakaratne, Falkner, and Atapattu 2019) , sometimes together with a few additional discoveries, e.g. in (Crichton et al. 2020; Pyysalo et al. 2019). Thus LBD evaluation has been mostly relying on the same small set of discoveries as benchmark for the past three decades. Consciously or not, LBD practitioners can be influenced by their knowledge of the target relations. This is a well-known methodological bias: in clinical trials, double blind experiments prevent both the patient and the researcher from influencing the outcome. For the same reasons, a solid LBD evaluation method would require a large and diverse set of target discoveries. In particular, the size of the test set is important in order to satisfy the condition of statistical representativity and consequently to ensure that the results can be generalized to other discoveries.

Poor LBD evaluation methodology might contribute to the lack of uptake by the biomedical research community at large. Despite a rich state of the art, LBD is still a mostly theoretical field. The lack of solid evaluation methodology is probably a factor which hinders the dissemination of LBD as a general research tool.

This is why we introduce the task of mining discoveries from the full existing literature. This task is very similar to LBD in the sense that both aim to produce relevant discoveries as output, but as opposed to LBD it does not have to predict *future* discoveries, i.e. it has access to the data after the time of discovery. By definition this makes the task easier since the system has access to more information. Yet the task is not trivial, because it involves filtering out a lot of relations which do not qualify as discoveries.

In order to formalize the concept of discovery we opt to focus on *impactful* discoveries, i.e. to use the impact of a relation in the literature as a marker of its discovery status: a significant discovery is expected to be followed by a surge in the number of mentions of the relation. We propose a method which calculates the trend across time for a relation based on its frequency in the literature. Significant surges are extracted, leading to a collection of impactful discoveries together with their time of impact. The results of the method are thoroughly analyzed and evaluated against a baseline representing the state of the art “time sliced” method. The resulting dataset is made available in two forms: the raw data can be used to evaluate or train LBD systems, and the visualization interface facilitates the manual exploration of the data.

The paper is organized as follows: the motivations and main idea are presented in §2. In §3 the method is described in detail, then the experimental results are analyzed in §4.

## 2 Approach

### 2.1 Motivations

Swanson 1986a,b introduced LBD as a method to explore “undiscovered public knowledge”, more precisely to identify missing links in a large and fragmented collection of knowledge. His approach, the ABC model, relies on the idea that different fields of specialization tend not to interact with each other. As a result, two subsets of the literature might each contain some knowledge about a shared concept, yet the lack of communication between the two fields can sometimes prevent potentially useful discoveries. In a broader sense, LBD can be defined as a task aimed at generating new research hypotheses, i.e. potential new discoveries Smalheiser 2012.

The field of LBD has seen significant progress since its inception. Kastrin and Hristovski 2021 systematically analyzed 35 years of LBD literature and observed that the field has grown in volume and diversity, developing from the initial text-based co-occurrences methods to advanced neural-based approaches (Crichton et al. 2020). However the question of the evaluation of LBD is still a major obstacle to its development as a mainstream research methodology. As many authors noted, e.g. (Ganiz, Pottenger, and Janneck 2005; Thilakaratne, Falkner, and Atapattu 2019; Yetisgen-Yildiz and Pratt 2009), evaluating LBD is very challenging due to the nature of the task: there is no direct way to assess whether the relations produced by a LBD system will eventually turn out to be significant discoveries.

Both Yetisgen-Yildiz and Pratt 2008 and (Thilakaratne, Falkner, and Atapattu 2019) provide a detailed review of the existing evaluation methods for LBD. The most commonly used is still by far the *replication* method: given a known discovery at time *t*, the LBD system is provided with the literature available before time *t* and produces a list (often a ranked list) of relations which represent potential “future” discoveries, i.e. from time *t* onward. The performance of the system is estimated based on how close to the top the target discovery is. While this evaluation method is reasonably sound, LBD systems are usually tested only against a small set of confirmed discoveries which have been previously found by existing LBD methods (traditionally the original discoveries made by Swanson 1986a, 1988). As already mentioned by Ganiz, Pottenger, and Janneck 2005, there are multiple biases in this evaluation methodology:

- There is a clear risk of confirmation bias, since the method is evaluated in terms of how well it retrieves discoveries that LBD methods are known to be good at finding.
- There is a risk of data leakage (including direct or indirect information from the test set into the model) when a new LBD system *A* is designed to improve over an older system *B* and both are evaluated on dataset *D*, especially if *D* consists of only a few instances. This can lead to overestimating the performance of system *A*.
- The performance obtained by evaluating on a small test set is not statistically reliable, and there is no way to measure the variance in performance. In other words, adding or removing an instance from the test set might drastically affect the overall performance. This makes any comparison between LBD methods very fragile.

Hristovski, Stare, et al. 2001 proposed a new evaluation method for their system, later formalized as a principled evaluation methology by Yetisgen-Yildiz and Pratt 2009 and called *time sliced evaluation* in (Thilakaratne, Falkner, and Atapattu 2019): given an arbitrary cut-off year *t* and a target term *x*, the cooccurrences of *x* which are found after time *t* but not before *t* are considered as gold standard discoveries. This approach solves most of the problems of the replication method, in particular it avoids any bias due to size or cherry-picking specific discoveries. As opposed to the replication method, it can also take into account positive and negative instances, making it possible to measure false negative cases in the LBD system output. This method was adopted for example by Lever et al. 2017.

The *time sliced evaluation* clearly solves the problem of using too few instances for evaluation, as almost every target term has multiple cooccurrences in the literature. However the use of the set of cooccurences as a proxy for the set of discoveries is a dramatic simplification. In fact, very few cooccurences represent a true discovery and the vast majority of the cooccurrences considered as “discoveries” for the purpose of this method are meaningless or poorly informative, for example when the cooccurrences appear by chance (e.g.*Ebolavirus* (D029043) and *Burnout, Professional* (D002055)) or when they involve at least one generic term (e.g. *Alzheimer’s Disease* (D000544) and *Elderly* (D000368)).

Thus with the *time sliced evaluation* method a LBD system is evaluated against a very large number of relations, but most of them are noise. While the true discoveries are included in the large amount of cooccurrences interpreted as gold standard, their proportion is unknown and likely low. As a result, the final performance does not reflect the ability of the system to predict insightful discoveries, only its ability to predict cooccurrences, which are often arbitrary. Despite solving serious methodological issues with the *replication* method, this method is still not fully satisfactory.

While LBD is unambiguously a data-driven task, there is no large-scale benchmark dataset available to evaluate and compare LBD methods. Clearly the field would benefit from such a resource, since it would bridge the gap between the replication method (actual discoveries but too few instances and only LBD-based) and time sliced evaluation (large number of examples but very noisy with respect to their discovery status). This also raises the difficult question of the definition of a discovery. Clearly there is no simple and objective definition, but at least some general criteria can be established. In this work we define a discovery as a meaningful relation between two concepts, i.e. a relation which satisfies these two conditions:

- There is evidence in the literature that the relation is valid (this excludes cooccurences which happen by chance);
- The relation must be insightful (i.e. relations which are trivially true are excluded).

We propose to interpret the concept of discovery as a spectrum which ranges from unambiguously positive cases to unambiguously negative cases. From this point of view, the few discoveries traditionally used by the *replication method* would belong to the small number of top positive cases, while the large set of cooccurrences used by the *time-sliced method* contains mostly negative cases. Our aim is to determine a measure of literature impact which reliably represents the “discovery level” of a relation across this spectrum. Such a measure would significantly improve the *time sliced evaluation* method, since it would allow selecting a subset of the most positive relations as discoveries, instead of the noisy full set of cooccurrences. In other words, our method can be seen as a filtering step applied to the gold-standard set of discoveries considered by the *time sliced* method: we propose substituting the large but low-quality gold-standard data with a smaller (but still large) number of high-quality relations in the *time sliced* evaluation method, preserving every other part of the method.^2^ Compared to the original *time sliced* method, this would lead to a much better separation between the positive and negative discovery cases, as illustrated in Fig. 1. With this modification, the *time sliced* evaluation would be more reliable thanks to the quality of the relations, and hopefully become more standard for LBD evaluation in the future.

**Figure 1:**
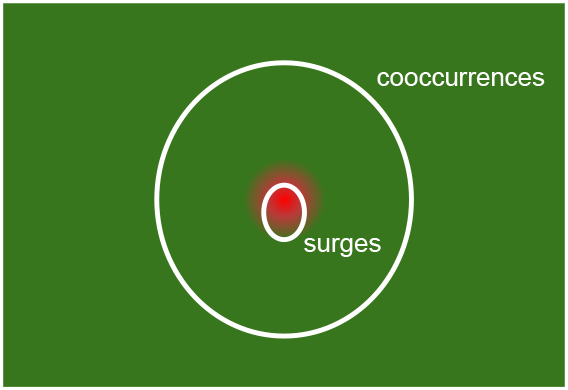
Schematic diagram of the positive/negative discovery cases. The full area represents the set of all possible relations, i.e. the cartesian product of the indvidual concepts. The red area represents relations which clearly qualify as discoveries (positive cases) and the green area represents relatons which clearly do not (negative cases). The gradient represents uncertain cases which might be considered on either side. The small white elipsis represents the target set of relations and the large white circle represents all the cooccurrences.

### 2.2 Impactful discoveries

In this article we propose a method to build a benchmark dataset of discoveries based on the impact that a discovery has on the scientific literature.^3^ The research ecosystem relies on community-based evaluation: through peer reviewing, journal reputation and citations, experts in a field evaluate each other’s work and estimate the value of their contributions. In this perspective, it seems intuitively sound to rely on the prevalence of a relation in the literature as an indicator for the interest or importance of this relation. In particular, one expects a significant discovery to generate a large increase in the number of mentions of this relations in the following years, as the community studies this discovery and explores its implications. It is reasonably straightforward to measure the frequency of any relation from a corpus of the target literature, for example using the collection of biomedical abstracts provided by Medline.^4^ Nonetheless the question of measuring *scientific impact* is not as simple as capturing a surge in frequency, because various factors other than a discovery event can affect frequency. Informally, the effect of a discovery on the literature can be described as a statistical outlier, i.e. an unusual event which stands out from the ordinary patterns.

It is worth noting that this approach assumes that the discovery is immediately recognized as such by the community and garners citations. Davies 1989 explains that some discoveries can be premature or postmature, both cases resulting in a lower impact (or no impact at all) in the scientific community. Assuming that literature impact is used as a proxy for discoveries, these false negative cases are possible but arguably rare.

Another potential limitation of this approach is that it is unclear whether impactful discoveries are the ones that matter for LBD. Some important discoveries might have a very low number of cooccurrences, for example because they happen in a small research community. In fact, it can be argued that rare relations are more likely to be overlooked, and therefore more likely to lead to a LBD-based discovery. In this sense, a system which performs well on a dataset of impactful discoveries might not be good at finding other types of discoveries. Nevertheless there is little doubt that impactful discoveries represent an important subset of discoveries, and could provide a decent proxy for the evaluation of LBD systems given the current limitations of the state of the art.

## 3 Method

### 3.1 Data and preprocessing

The experiments carried out in this paper use Medline as the source literature. Medline is a database containing more than 31 millions references to articles published in life sciences journals.^5^ Every reference in Medline is annotated with a set of MeSH descriptors^6^ which represent the main biomedical concepts relevant to the article. We opt to use the Medline MeSH decriptors as concepts because these have been carefully annotated/reviewed by experts, thus the risk of error in the data is very low. However MeSH concepts are coarse and generic compared to other biomedical ontologies. The method described below can be applied to alternative representations of the literature, for example using PubTatorCentral^7^ (Wei et al. 2019) or the UMLS Metathesaurus^8^. It could also be applied to richer semantic descriptions of the relations, e.g. DrugX-TREATS-DiseaseY, such as SemRep^9^^10^

The year of publication and the MeSH descriptors are extracted for every entry using a modified version^11^ of the “Knowledge Discovery” code by Lever et al. 2017. Every pair of MeSH descriptors in the same article defines a cooccurrence between two concepts. The raw data is processed using the “TDC Tools” repository^12^ in order to obtain the frequency of (1) every concept (MeSH descriptor) by year and (2) every cooccurrence between two concepts by year. Additionally the pairs with less than 100 cooccurrences across all years are removed in order to prevent a large amount of noisy relations in the data.^13^ The range of years is also filtered from 1950 onwards to avoid the low data volume of the early years.

Most of the examples and experiments presented below rely on a subset of the literature related to Neurodegenerative Diseases (NDs). This subset is obtained by first selecting all diseases which have the concept *Neurodegenerative Diseases* (D019636) as ancestor in the MeSH hierarchy, and then selecting all the concepts which cooccur at least once with any of these target concepts.^14^ The final dataset contains 291k distinct relations and 1,8k distinct concepts (see details in Table 1) with their frequency by year.

**Table 1:** Number of concepts and relations at different stages of filtering.

| Data | unique concepts | unique relations |
| --- | --- | --- |
| Full data | 29,406 | 47,291,526 |
| Frequency $\geq 100$ | 26,268 | 1,813,710 |
| ND filtered | 1,803 | 290,797 |

### 3.2 Measuring literature impact

In order to detect literature impact, a *trend* indicator is calculated for every year *y* and every relation (*c*_1_, *c*_2_). First, a moving average over a window of *n* years is applied in order to smoothen the frequency variations and capture the trend more acurately. Then a measure of statistical association is calculated for every year independently. Similarly to other LBD works (Crichton et al. 2020; Pyysalo et al. 2019), we use a set of standard measures: Pointwise Mutual Information (PMI) as well as Normalized PMI (NPMI), the latter being less biased than PMI towards low frequency relations Bouma 2009; Mutual Information (MI) and Normalized MI, which take into account all the cases of concepts *c*_1_ and *c*_2_ appearing or not (we follow the definitions from (Bouma 2009)); and Symmetric conditional probability (SCP), the product of the two conditional probabilities. Although the joint probability is unlikely to be a good indicator, it is included as a baseline measure.^15^

Figure 2 shows the evolution between 1988 and 2018 of the joint probability and other measures between *Amyotrophic Lateral Sclerosis* (ALS) and five distinct concepts. These cases illustrate the diversity of the patterns captured by different measures. For example, the relations of ALS with concept *house mice* or *middle age* have important changes across time in terms of probability (first row), but they do not, or less, in terms of statistical association (remaining rows). On the contrary, *C9orf72 Protein*^16^ and *riluzole*^17^ have moderate frequency changes but show a strong surge according to some of the association measures. The relation of ALS with *anterior horn cells*^18^ has a moderate evolution across time according to most measures.

**Figure 2:**
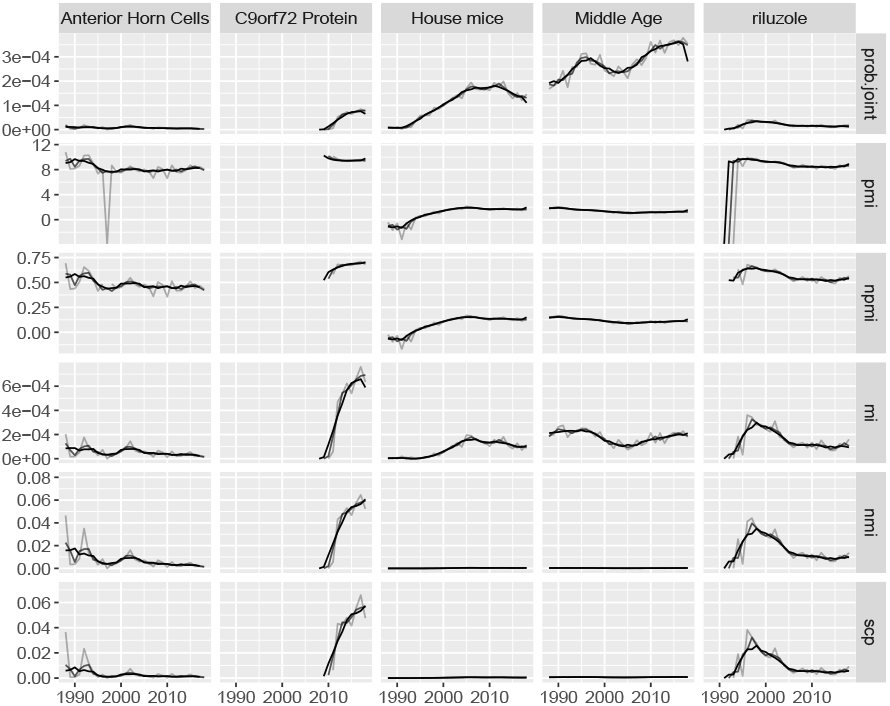
Joint probability and other association measures in the period 1988-2018 for the concept *Amyotrophic Lateral Sclerosis* (D000690) paired with five concepts: *Anterior Horn Cells* (D000870), *C9orf72 Protein* (D000073885), *House mice* (D051379), *Middle Age* (D008875) and *Riluzole* (D019782). In one plot the different curves represent the values for different moving average window sizes, from the smallest size 1 (lightest) to the highest size 5 (darkest).

The association measure is meant to represent the importance of a relation at any given time. The next step consists in measuring how the importance of the relation evolves across time, i.e. its *trend* from one year to the next. Two simple indicators are considered:

- *diff* is the difference between the association values of the two years: *v_y_* − *v*_*y*−1_.
- *rate* is the relative rate of the association values of the two years: 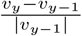.

### 3.3 Detecting surges

Since the trend indicator is a continuous numerical value, a threshold parameter is needed in order to separate regular years (negative, low or moderate trend value) from the years marked by a surge (high trend value). Naturally, the level at which a trend indicator becomes a surge is subjective and may depend on the target application.

In the case of building a test set of discoveries for LBD evaluation, the cases of interest are the most dramatic surges since one wants to maximize precision, i.e. to minimize false positive errors. Figure 3 shows two views of the trend distribution for the NMI measure with the *diff* indicator (other measures and indicators show the same general pattern). Even on a logarithmic scale, the regular histogram is not very informative due to the extremely high proportion of points close to zero. The quantile plot is more insightful, as it shows two clear inflection points at the extreme ends of the curve: while the vast majority of the points lie so close to zero that the curve looks flat, at both ends a few points get significantly farther from zero. The inflection point is determined by finding the point which maximizes the product of the normalized trend and the quantile (i.e. finding the largest rectangle starting from the top left corner of the quantile plot and having its bottom right corner on the curve).

**Figure 3:**
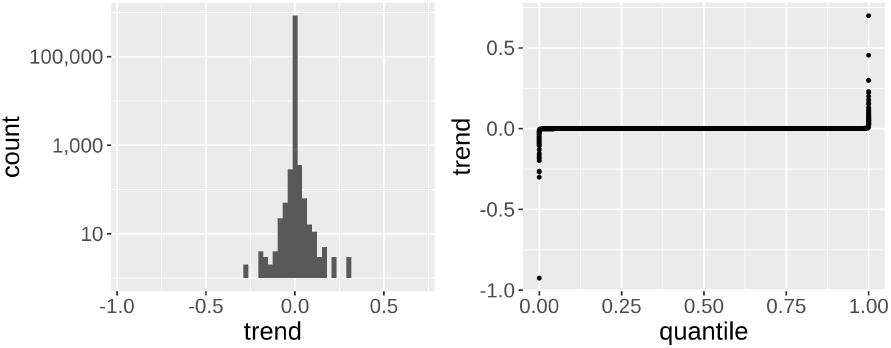
Two views of the trend distribution for 20,000 random relations (NMI, window size 5, diff indicator). Left: histogram with logarithmic scale on the Y axis. Right: quantile plot (data points sorted by trend value on the X axis with their value shown on the Y axis).

This method leads to selecting only the cases where the trend is exceptionally high, thus obtaining a very small proportion of pairs (relation, year) above the cut-off point: among the combinations of 6 measures, 2 trend indicators and 3 sliding window sizes (1, 3 and 5), the median number of surges found represents only 0.2% of the pairs (relation, year) and 2.9% of the unique relations (cooccurrences). The majority of the combinations of parameters results in between 800 and 30,000 unique relations with surges, among a total of 290,797 possible relations (between 0.2% and 10%).

### 3.4 Post-processing

Several optional post-processing steps are also implemented, their use depends on the target application:

- The discovery year can be adjusted in cases where the surge is found in a year where the real frequency is zero. This may happen due to the moving average window, which occasionally creates high frequency values before the relation even exists. In such cases the surge year is shifted to the next year with non-zero frequency within the moving average window. Additionally, in the perspective of mining discoveries, the earliest surge is selected whenever a relation is found to have several years with a surge (i.e. any surge year later on is filtered out). However multiple surges in a relation can be meaningful in the context of some different application, for instance in the exploration of the relations which are “trending” at some particular point in time.
- Initial observations show that a large number of relations involve abstract and/or generic concepts, such as *Data Interpretation, Statistical* (D003627), *Cross-Over Studies* (D018592) or *Quality of life* (D011788). These relations may reflect real evolutions of the biomedical domain, especially methodological and technical innovation in biomedical research, but they are not primarily biomedical in nature and are often difficult to date precisely. This is why the UMLS Semantic Groups^19^ can be used to filter the concepts types, by default selecting only the relations involving the four groups *Anatomy*, *Chemicals and Drugs, Disorders* and *Genes & Molecular Sequences*.
- A significant number of relations involve two concepts very closely related to each other, such as *Synucleins* (D051843) and *alpha-Synuclein* (D051844), *NF2 gene* (D016515) and *Neurofibromin 2* (D025581), *Presenilin-1* (D053764) and *Presenilin-2* (D053766). Naturally these relations tend to have simultaneous surges when their main concept has a surge within another relation since the two related concepts are frequently mentioned together. These spurious cases can also be filtered out by discarding relations in which at least one of the concepts has a high conditional probability with the other, indicating a trivial relation. The selection of the threshold is a trade-off between precision and recall: a high threshold means that the resulting list includes some trivial relations, whereas a low threshold eliminates some real discoveries from the list.

## 4 Results and analysis

### 4.1 Evaluation

Naturally, evaluating the task of mining (past) discoveries faces the same issue as LBD: the concept of discovery is interpretative and there is no large-scale dataset of discoveries available (see §2.1). However the former task has the advantage that its “predicted” discoveries can be qualitatively assessed, since their significance in the field is known, at least by experts. The evaluation of our method is decomposed into two main steps:

First, we analyze different aspects of the method using the ND subset: impact of the parameters, accuracy of the predicted year of discovery, qualitative analysis. A gold standard dataset of 12 important ND discoveries is created in order to quantitatively analyze the method. This dataset is built using external sources to collect the year of discovery.^20^ The choice of the relations is arbitrary and is partly based on whether the relation has a clearly established year of discovery, therefore the dataset cannot be interpreted as a representative sample. Even in the case of discoveries which are clearly recognized in the ND field, there are often ambiguities about the exact date.^21^ x This ND gold standard dataset is used to measure recall, i.e. the proportion of gold discoveries detected by the method. A discovery is considered as detected if the relation is retrieved and its surge year is within the window *y* ± *n*, where *y* is the true discovery year and *n* is a fixed constant.

Second, we compare our method against a baseline representing the original time sliced method. For our method we select specific values for the parameters, based on the previous analysis (*SCP/diff/5*, see below). The baseline set of discoveries is obtained by extracting the *N* most frequent relations in the full dataset, together with their first year of cooccurence (see details in Table 1).^22^ *N* is chosen to be equal to the number of relations returned by our method (*N* = 9, 092), so that the two list of relations are comparable. The same post-processing steps are applied to both methods: filtering years 1990-2020, maximum conditional probability 0.6, and filtering of the four groups *Anatomy, Chemicals and Drugs, Disorders* and *Genes*. The two resulting lists of discoveries are evaluated as follows: for each list, the top 100 relations are selected as well as a subset of 100 relations picked randomly in the list. Then the four subsets of 100 relations each are randomly shuffled into a large dataset which is then annotated manually. The final list contains no indication of which subset a relation comes from, so that the annotator cannot be influenced in any way. The annotation process is simplified in order to minimize the subjectivity involved in deciding whether a relation qualifies as a discovery or not. Every pair of concepts is labelled as one of three possibilities ‘yes’, ‘no’, ‘maybe’ regarding the discovery status. The annotator relies on Google Scholar queries with the two concept terms in order to determine their status:

- If the top results show some evidence of a significant, non-trivial, new and impactful relation between the two concepts, then the relation is annotated as ‘yes’. This requires at least one fairly clear title or abstract mentioning the relation as a discovery, with a healthy number of citations. The year of the main article is reported as gold-standard year (whether it is close to the predicted year or not).
- If there is evidence that the relation is either trivial, questionable or has very little impact (few papers or citations), then it is labelled as ‘no’. This includes for example obvious relations (e.g. “Adrenergic Receptor - Adrenergic Antagonists”) and too trivial terms (e.g. “Traumatic Brain Injury - Neurons”).
- In any other case, the status is considered ambiguous and the relation is labelled as ‘maybe’. This includes cases where the annotator cannot understand the articles, has doubts about the originality, or the citation count is moderate. These cases are ignored in the evaluation results.

It is worth noting that this annotation policy is fairly strict regarding the discovery status of the relation: for example, in many cases there is a discovery with one of the terms but the other term is only indirectly related; such cases would be labelled as negative. The non-trivial condition also discards many relations which could potentially qualify as discoveries. These strict criteria are intended to make the annotation process as deterministic as possible, but real applications of LBD might consider a larger proportion of predicted discoveries as relevant. Thanks to this annotated dataset, it is possible to estimate the precision (proportion of true discoveries among the predicted ones) of our method and compare it against the baseline.

### 4.2 Parameters

The effect of the different parameters of the method (see §3.2) is evaluated by comparing their performance against the ND gold standard dataset. The surges are extracted for every configuration of parameters among:

- Three window sizes for the moving average (1,3 and 5);
- The six association measures (joint probability, PMI, NPMI, MI, NMI and SCP)
- The two trend indicators (*diff* and *rate*)

For every parameter, figure 4 shows the average performance (recall) on the dataset, averaging across the values of the other two parameters. Performance increases with the window size of the moving average: 5 is better than 3, which is better than no moving average (1). The NMI and SCP measures perform best, followed by NPMI. PMI and MI perform poorly, more so than even joint probability. Finally the *diff* indicator performs drastically better than *rate*. Importantly, the *first year* filter (see §3.3) decreases performance only slightly compared to keeping all the surges; this is an indication that the method works as intended: for every relation which has several high surges, the first surge is very likely to be detected around the true time of discovery.

**Figure 4:**
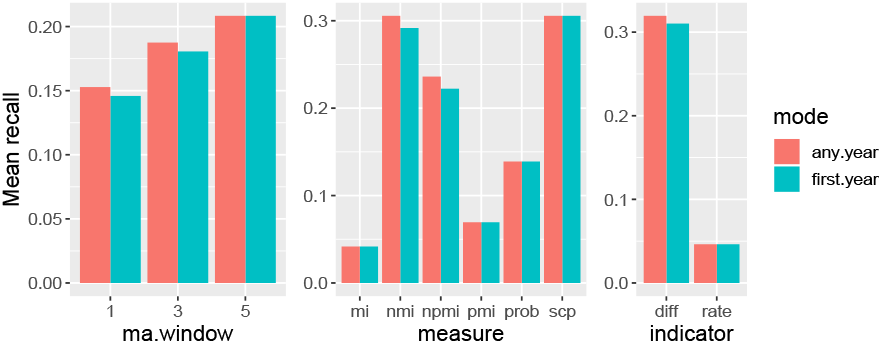
Average recall within *y* ± 3 years on the gold standard ND dataset (12 discoveries) by parameter. For each parameter value, the mean recall is calculated across the values of the other two parameters. The two colours represent whether all the surges are taken into account (*any year*) or the first year filter is applied (*first year*, see §3.3).

The best performing individual configurations are consistent with the results by parameters. SCP/diff/5 performs best, identifying the correct discovery year in 8 cases out of 12 (recall 0.67). It is followed by 6 configurations which perform equally well (7 cases out of 12; recall 0.58): NMI with window 1, 3 or 5, SCP with window 1 or 3 and NPMI with window 5 (all with the *diff* indicator). These results confirm the superiority of the NMI and SCP measures, but the small size of the dataset does not allow any fine-grained comparison between the configurations.

An analysis of the correlation between the results of the different configurations is also carried out in order to gain insight about the parameters independently from the gold standard dataset (the *rate* indicator is discarded due to its very poor performance). The level of agreement between two configurations is measured as the proportion of relations in common (overlap coefficient), considering that a relation is in common between two configurations if (1) it has a surge in both and (2) the two detected surges are within 5 years of each other. Figure 5 shows that the configurations tend to have high overlap values in general, considering the small proportion of relations and the close surge year constraint. As expected, the highest similarity values are obtained within the same measure, varying only the window size. But similarity is also high between different measures: NMI and SCP share 70 to 80% of their surges, while PMI and NPMI share more than 80% but only using the same window size. These results are consistent with the results obtained on the gold dataset, adding to the evidence that the NMI and SCP measures return surges characteristic of a discovery pattern.

**Figure 5:**
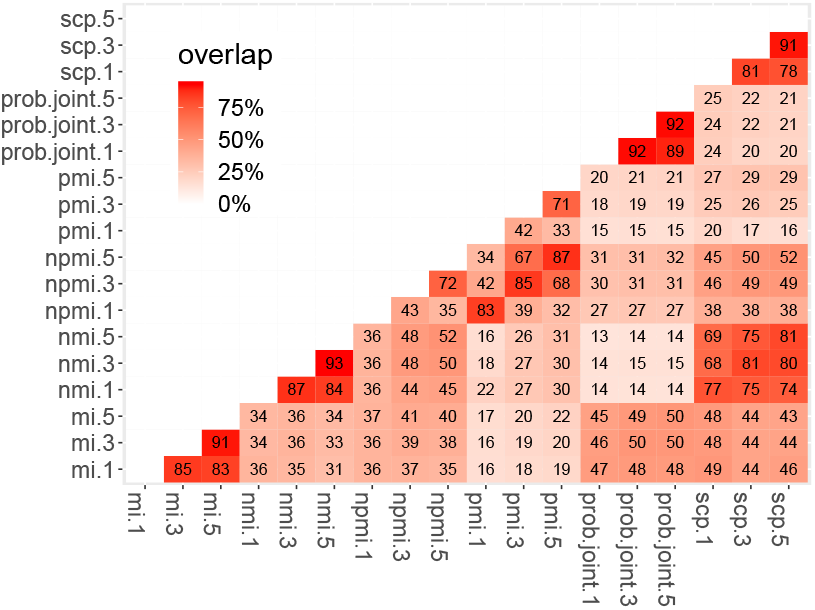
Overlap coefficient for every pair of configurations of parameters (*diff* indicator). A relation is considered in common between two configurations if their corresponding surges are within 5 years of each other.

### 4.3 Discoveries across time

In this part we investigate the distribution of the surges across time. There are various potential biases related to the time dimension: for example, the volume of data is not uniform across time, since the number of entries in Medline increases by approximately 4% every year. There can also be artifacts due to the construction of Medline as a resource.^23^ A distribution of the detected surges can be observed in figure 6 (middle); the distribution of the first cooccurrence year for all relations, i.e. the input data, is also shown for comparison (top). The surges patterns follow the first cooccurrence patterns quite closely, evidencing the absence of any visible bias due to the surge detection method. Most of the years have between 300 and 600 surges, which represents around 10% of the number of first cooccurrence relations. The peak in the 1960s is likely an artifact due to the construction of Medline. The decrease after the 2000s may be partly due to the filtering of the relations which have less than 100 cooccurrences (see §3.1), since relations which appeared in recent years had less time than older ones to accumulate 100 mentions. It is possible that the early years (50s and 60s) contain some spurious surges due to the low volume of data and the introduction in the data of concepts which might have existed before. For example, the relation *Adrenal Medulla* (D000313) and *Pheochromocytoma* (D010673) is detected as surging in 1952, although the discovery happened earlier.Nevertheless other cases among the earliest surges detected appear to be valid, such as the relation between *adenosine triphosphate* (D000255) and *Stem cells* (D013234) which has a surge detected in 1952 (Bendall 1952a,b).

**Figure 6:**
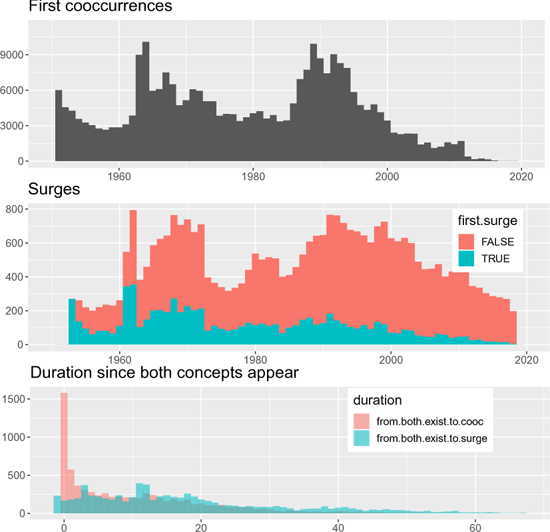
Top: number of unique relations with frequency higher than 100 by year of first cooccurence. Middle: number of surges by year, in blue for the earliest surge of a relation, red for subsequent surges. Bottom: difference between the earliest year where both concepts appear individually and (1) their first cooccurrence (red), and (2) their first detected surge (blue). Surges parameters: SCP measure, window 5, *diff* indicator.

We also study how long after the introduction of the two concepts surges happen. Given a relation between two concepts *c*_1_ and *c*_2_ which appear for the first time at years *y*_1_ and *y*_2_ respectively, the earliest possible year for a cooccurrence (and consequently for a surge) is *max*(*y*_1_, *y*_2_). The bottom plot of figure 6 shows the distribution of (1) *y_c_* − *max*(*y*_1_, *y*_2_) and (2) *y_s_* − *max*(*y*_1_, *y*_2_), with *y_c_* the year of the first cooccurrence and *y_s_* the year where the first surge is detected. With parameters *SCP/diff/5*, the first surge happens in average 7.6 years after the first cooccurrence. Among the relations which have a surge, 21% have their first cooccurrence the same year as the two concepts appear, whereas only 5.3% have their first surge this year. Similarly, 10% of the relations have a surge in the first two years while 30% of cooccur in the first two years. Thus in a large number of cases, the first cooccurrence appears at the same time or soon after the first year where both concepts exist. However the first surge appears later most of the time, indirectly illustrating an important difference between the time sliced method and our method: the former confuses cooccurrence and discovery, whereas the latter waits for evidence that the relation is an actual discovery.

In most case the first surge occurs within 20 years of the two concepts appearing in the data. Nevertheless it is also possible for the surge to happen much later; in some cases, both concepts exist from the starting point of our dataset (1950) and have a first surge in the 2010s. For example, *Spinal Muscular Atrophy* (SMA; D009134) and *Oligonucleotides* (D009841) first appear in the data in 1951 and surge only in 2016, when a clinical trial for an antisense oligonucleotide drug for SMA proved successful (Finkel et al. 2016). However there are also questionable cases with some relations between two general concepts; for example, the relation between *Muscle Spasticity* (D009128) and *Motor Neuron Disease* (D016472) has its first detected surge in 2017, while both concepts exist in the data since 1950. This case may correspond to a “gap”, as described by Peng, Bonifield, and Smalheiser 2017.

### 4.4 Comparison to time sliced baseline

As explained in 4.1, our method is compared against a baseline set of relations ordered by frequency, representing the time sliced method. The original time sliced method would normally include every cooccurrence which appear in the data: in our dataset (after relations with less than 100 cooccurrences were filtered out), this represents 108,794 unique relations after applying the post-processing steps, compared to 9,092 for our method with the parameters *SCP/diff/5*.

In the results presented in table 2, only the 9,092 most frequent relations are considered for the base-line in order to compare the two methods on a fair basis. Surprisingly, the subset made of the top 100 relations (by frequency for the baseline, by trend for our method) obtains a lower precision than the subset made of 100 random relations, even though the top relations are expected to be more likely discoveries.^24^ On both subset the difference in precision between our method and the baseline is quite large (14.5 and 20.5 points), but the *χ*^2^ test between the ‘yes’ and ‘no’ categories is significant only for the top 100 subset.

**Table 2:**
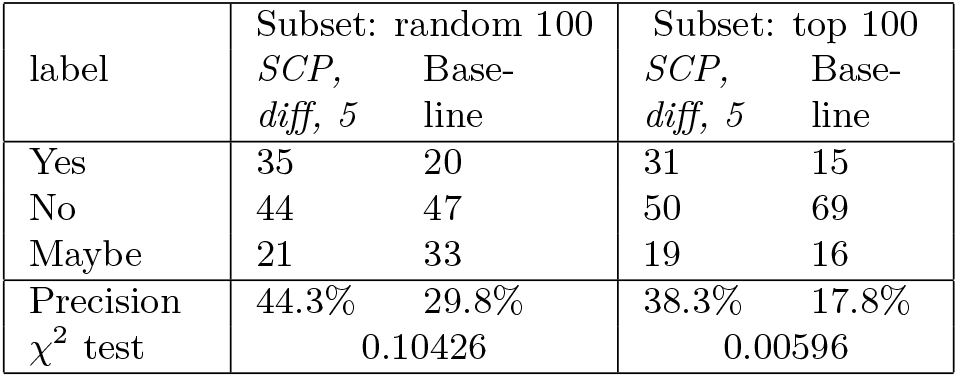
Results of the SCP surges vs. *time-sliced* baseline on 400 manually annotated relations. The data was divided into 2 subsets for both methods: random selection of 100 relations (among the full 9092 relations), and top 100 relations (see details in §4.1). The p-value of an *χ*^2^ test between the yes/no cases is shown.

## 5 Discussion

### 5.1 Advantage for LBD evaluation

As explained in §2.1, the *time sliced* method proposed by Hristovski, Stare, et al. 2001 and Yetisgen-Yildiz and Pratt 2009 is the only existing formal evaluation paradigm for LBD, and is often not even used. The proposed method improves the *time sliced* method by filtering out a very large proportion of mostly negative cases. As shown in §4.4, our method obtains a higher precision than the baseline, considering that this comparison does not take into the 92% least frequent relations of the *time sliced* method. Our method uses literature impact as a marker for “true” discoveries, leading to considering only a very small subset of the cooccurrences as positive cases: in the experiment presented above, the median number of relations above the surge threshold represents 3% of all the input relations (see §3.3). Thus our method allows a more accurate balance between the positive and negative cases:

- In the *time sliced* method, the cases considered as negative are perfect (since there cannot be a discovery without cooccurrence), but the vast majority of the cases considered as positive are actually negative.
- In our method, both the cases considered as negative and positive are slightly imperfect, but the latter are likely to represent true discoveries.

### 5.2 LBD and the time dimension

Traditionally, LBD models consider a static representation of the state of knowledge at a particular point in time. The time dimension is only taken into account for the purpose of evaluation. This implies that the existing relations in the dataset are considered equivalently likely to lead to a future discovery. Clearly, this is also a simplication: the scientific knowledge ecosystem follows a dynamic evolution, with some topics trending and others being abandoned (see e.g. Kastrin and Hristovski 2019). The identification of a significant new relation often causes a thrust of research, simply because the new relation is more likely to help uncover more new knowledge than any older relation. It seems intuitively relevant for LBD methods to integrate the time dimension as well. For example, the ABC model relies on linking concepts in common between two heterogeneous subsets of the literature and drawing new relations by transitivity; it might be relevant as well to prioritize relations which were discovered at different times. More advanced LBD models might even be able to model the dynamic evolution of the topics and concepts across time in order to actually predict future trends, instead of only exploring the search space of all possible relations.

### 5.3 Exploration tool and other applications

As a result of this work, two datasets of surges detected from Medline are made available: the ND subset studied in this paper and the surges obtained on the full Medline data.^25^ A simple exploration tool is also made available.^26^ This tool lets a user observe the relations retrieved by our method with various filtering options. For example, a user may search for recent discoveries related to treating a particular disorder by selecting the appropriate target concept, range of years and semantic group. This kind of tool offers a very synthetic view on the body of knowledge represented in the biomedical literature. More generally, the availability of a dataset of time-stamped discoveries may lead to novel applications and usages, for example as a methodological resource for systematic reviews.

## 6 Conclusion

This work demonstrates that it is possible to exploit the evolution of the literature across time in order to retrieve the relations which had the most significant impact in the past. It also provides evidence that the surges extracted through this method correspond to impactful discoveries, for the most part. Compared to previous work in the evaluation of LBD, this approach can be used to replace the set of cooccurrences used as gold standard by the time sliced method, hence offering a more meaningful way to evaluate LBD systems on a large set of discoveries. It might also pave the way for more fine-grained LBD methods, which could exploit these past discoveries to train supervised models.

Although simpler than LBD, the task of extracting impactful discoveries from the existing literature is not trivial. To some extent, the ability to reliably identify past discoveries can be seen as a pre-requisite for the harder case of LBD, since the latter also needs to identify discoveries but with less information available as input. In particular, the distinction between meaningful and non-meaningful relations is complex. Existing LBD methods rely on simplifications which tend to mask this difficulty, but this question must be addressed if LBD is to become a well grounded methodological tool for the biomedical domain.

## Acknowledgements

The authors are grateful to the three anonymous reviewers for their valuable comments. Erwan Moreau also thanks Ashjan Alsulaimani for her suggestions.

## Footnotes

1 We use the word *relation* throughout this work for any (potential) relationship between two concepts. In general the nature of the relationship is unknown; in theory the association between the concepts it does not even have to be positive.

2 In particular, one should still use the first year of cooccurence as the cut-off year in order to avoid any data leakage.

3 It is worth noting that alternative methods could be considered to mine discoveries, for example manual curation by experts, mining review articles, based on bibliometrics, etc.

4 https://www.nlm.nih.gov/databases/download/pubmed_medline.html

5 The version of the data used in the experiments was downloaded in January 2021. The full code and instructions are provided at https://github.com/erwanm/medline-discoveries.

6 https://meshb.nlm.nih.gov

7 https://www.ncbi.nlm.nih.gov/research/pubtator/

8 https://www.nlm.nih.gov/research/umls/index.html

9 https://lhncbc.nlm.nih.gov/ii/tools/SemRep_SemMedDB_SKR/SemRep.html.

10 Naturally such rich semantic relations are very useful for LBD Hristovski, Friedman, et al. 2006. But the extraction of the relations from the text is a potential source of error, this is why we adopted the most basic and reliable representation available.

11 https://github.com/erwanm/knowledgediscovery.

12 https://github.com/erwanm/tdc-tools

13 Rationale: relations with less than 100 occurrences are less likely to have an exceptionally high impact, i.e. to qualify as “impactful discoveries”.

14 This is equivalent to filtering all the articles which contain at least one of the target concepts and then selecting all the concepts in these documents. This filtering is purposefully loose in order for the dataset to include a broad range of concepts and relations with varying levels of specificity.

15 As opposed to using the raw frequency, the probability prevents the increase in number of publications (around 4% more every year) to artificially inflate the trend of a relation.

16 The identification in 2009 of C9orf72 repeat expansions as the most common genetic variant associated with both ALS and frontotemporal dementia (FTD) confirmed the previously recognized pathobiologic association between the two conditions.

17 *Riluzole* is a drug used in the treatment of ALS; it was introduced in the US in 1995 and in the EU in 1996.

18 A nerve cell in the spinal cord, rhombencephalon, or mesencephalon.

19 https://lhncbc.nlm.nih.gov/ii/tools/MetaMap/documentation/SemanticTypesAndGroups.html.

20 This criterion was established to ensure that the relations were uncontested discoveries. It turned out that this caused the rejection of many candidate relations, leaving only a few discoveries. Nonetheless it was decided not to relax the conditions, i.e. to favour quality over quantity.

21 E.g. the causal gene for Huntington’s disease was approximately located in 1983 but precisely located on chromosome 4 in 1993.

22 We are grateful to an anonymous reviewer for suggesting this idea. The ranking by frequency is commonly used in LBD systems in order to show the most important relations first, therefore it is a reasonable baseline method.

23 For example, the proportion of entries with an abstract in Medline jumps from 5% in 1974 to 40% in 1975

24 This could be due in part to the high randomness and possible annotation errors, but in the case of the baseline many of top relations are visible outliers: due to the particular severity of the pandemic, the most frequent relations involve *Covid19* with various other generic concepts, often not qualifying as a discovery (e.g. *Coronavirus* and *Emotional stress*).

25 https://zenodo.org/record/5888572.

26 https://brainmend.adaptcentre.ie/.

